# Progenitor Heterogeneity in the Developing Cortex: Divergent and Complementary Roles of NG2-Progenitors and RGCs

**DOI:** 10.1101/2025.01.27.635022

**Authors:** Ana Cristina Ojalvo-Sanz, María Figueres-Oñate, Sonsoles Barriola, Carolina Pernia-Solanilla, Rebeca Sanchez-Gonzalez, Lina Delgado García, Laura López-Mascaraque

## Abstract

Brain development is a highly coordinated process that arises from a pool of neural progenitor cells (NPCs). Traditionally, research has largely focused on Radial Glial Cells (RGCs), which produce neurons and glia. However, recent findings revealed that the progenitor landscape is more complex and heterogeneous than previously understood. Emerging evidence indicates that NG2 glia, also known as oligodendrocyte precursor cells (OPCs), may contribute to generating other cell types, including neurons and astrocytes. This suggests that NG2 glia, or a specific subset, may act as multipotent NPCs, expanding their role in brain development beyond their known lineage.

We explore the cellular, molecular, and functional differences between NG2- and GFAP- expressing progenitors (NG2-NPCs and GFAP-NPCs) across different developmental stages. Using *in-utero* electroporation and StarTrack technology, we examined whether these progenitors represent distinct populations and analyzed how their differences influence their functional roles based on cell fate decisions. Our findings uncover functional divergence between NG2-NPCs and GFAP-NPCs, particularly regarding their cell fate decisions. While both populations contribute to the formation of neurons and glial cells, the progeny of NG2-NPCs and GFAP-NPCs differ markedly in their contributions to neurogenesis and gliogenesis. Further, we conducted a comprehensive transcriptomic analysis of NG2-NPCs and GFAP-NPCs to elucidate the molecular basis for these functional differences. This study demonstrates distinct gene expression profiles. While NG2-NPCs show enrichment for genes associated with neurogenesis and synaptic transport, GFAP-NPCs displayed upregulation of genes involved in progenitor cell maintenance. This indicates that each progenitor type is governed by distinct molecular programs, shaping their contributions to brain architecture and function. In summary, our findings underscore the divergent roles of NG2-NPCs and GFAP-NPCs in driving cellular diversity and functional maturation in brain development, providing new insights into how these progenitor populations contribute to neural diversity and brain function.

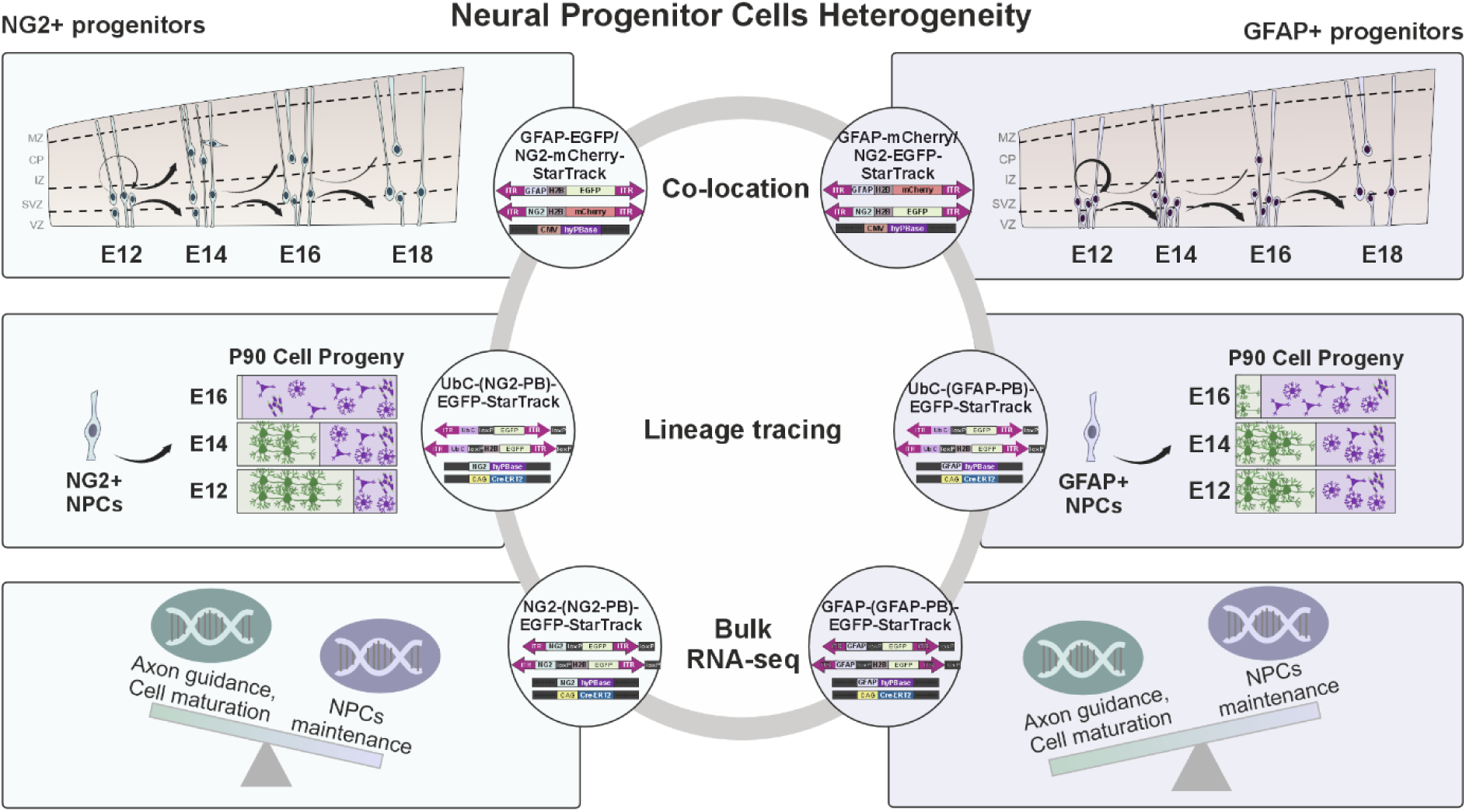

## 1. Introduction

Brain development is a highly orchestrated process that depends on the precise function and coordination of neural progenitor cells (NPCs). These cells give rise to a diverse array of neuronal and glial cell types, which together construct the intricate architecture of the central nervous system (CNS) (1–6). Traditionally, research has primarily focused on Radial Glial Cells (RGCs), long recognized as the principal NPCs during embryonic development, responsible for generating both neurons and glia (1–7). However, recent advances in transcriptomics and lineage-tracing technologies have uncovered a far more complex progenitor landscape within the developing brain, marked by extensive heterogeneity. A growing body of evidence reveals that NPCs within the brain are not a homogeneous population but instead comprise distinct molecularly and functionally diverse subsets, each with unique temporal and spatial roles in neurogenesis and gliogenesis (8–19). This diversity is crucial for ensuring the precise patterning of brain development, as different progenitor populations contribute to specific cell types and shape the complex architecture of the CNS. These findings highlight the importance of understanding the heterogeneity of neural progenitors to fully comprehend how the brain is formed and how it may go awry in neurodevelopmental diseases.

One cell type that has recently garnered significant attention is NG2-glia, also known as oligodendrocyte precursor cells (OPCs). These cells are characterized by their expression of the nerve/glial antigen 2 (NG2) proteoglycan, encoded by the *Cspg4* gene (20,21). Traditionally, NG2-glia have been associated with oligodendrogenesis, playing a role in forming oligodendrocytes during development (21–24). However, evidence suggests that adult NG2-glia may contribute to generating other cell types, such as neurons and astrocytes, under certain conditions (25–32). While NG2-glia are well established in postnatal gliogenesis, their roles and presence during early development remain poorly understood. It has been proposed that a subset of NG2+ cells could act as progenitors during critical stages of brain development (33). Nevertheless, it remains unclear whether NG2-expressing neural progenitor cells (NG2-NPCs) represent a distinct population during early brain development or how they relate to the well-studied RGCs. Specifically, it is unknown if NG2-NPCs and RGCs act as separate progenitor pools with unique developmental trajectories or overlap in location and molecular identity. Clarifying these distinctions is crucial, as it could shed light on brain development and the mechanisms driving cellular diversity in the CNS, which are fundamental to understanding numerous neurodevelopmental disorders (34–36).

This study provides an in-depth analysis of cellular and molecular diversity among NPCs, focusing on the role of NG2-NPCs in early brain development. We use genetic tools to co-localize NG2-NPCs with other progenitor populations, such as RGCs. Through lineage-tracing techniques, we map the developmental trajectories and specific roles of NG2-NPCs relative to RGCs. Finally, transcriptomic analyses reveal distinct molecular signatures that further differentiate NG2-NPCs from RGCs. Together, these findings provide new insights into how the heterogeneity of NPCs shapes brain development, offering a deeper understanding of the origins of cellular diversity in the CNS and how this diversity contributes to neural network formation and function.

## 2. Material & Methods

### 2.1. Mouse line

Adult pregnant wild-type C57BL/6 mice were obtained from the Cajal Institute (Madrid, Spain) and housed in the institution-accredited animal facility. All experimental procedures were conducted following the ethical guidelines established by the European Union (Directive 2010/63/EU) and the Spanish Ministry of Science and Innovation (RD 1201/2005 and L 32/2007). The study protocol received approval from the Bioethical Committees of the Cajal Institute, CSIC, and the Community of Madrid (PROEX 314/19). Mice were maintained on a 12-hour light-dark cycle with a room temperature of 22°C. Embryonic day (E0) was determined by the detection of a vaginal plug, while the day of birth was designated as postnatal day 0 (P0). A minimum of N=3 animals per group was used for all experimental procedures to ensure statistical robustness.

### 2.2. StarTrack Modifications for Specific Progenitor Lineages

To tag NG2-progenitors (NG2-NPCs), GFAP-progenitors (GFAP-NPCs), and their progenies, we employed the StarTrack method, a multicolor genetic tool to generate a unique, inheritable “color-code” within progenitor cells and their offspring (10,37). This method uses the PiggyBac transposon system (38), integrating plasmids that codify fluorescent proteins (XFPs) into TTAA sites in a stable and heritable manner (10,37).

For progenitor-specific tagging, we used ubiquitous PiggyBac plasmids under the control of lineage-specific transposase promoters. Specifically, the *NG2* promoter (NG2-hyPBase), provided by Prof. Sellers, was used to target NG2-NPCs, while the human *GFAP* promoter (GFAP-hyPBase), from Dr. Lundberg, was employed to label GFAP-NPCs. GFAP-NPCs. Plasmid amplification and purification were performed using the NucleoBond Xtra Maxi kit (Macherey-Nagel; France #740414.50).

We combined PiggyBac plasmids driven by the NG2 or GFAP promoters with the ubiquitous CMV-hyPBase transposase for spatial colocalization experiments. In progeny assessments, UbC-EGFP + UbC-H2B-EGFP plasmids were co-electroporated with GFAP-hyPBase or NG2-HyPBase and Cre-ERT2 to monitor lineage-specific differentiation. Transcriptomic analyses were conducted using plasmids and transposases controlled by the same promoter, such as NG2-EGFP + NG2-H2B-EGFP with NG2-hyPBase or GFAP-EGFP + GFAP-H2B-EGFP with GFAP-hyPBase.

### 2.3. *In utero* electroporations (IUE)

*In utero* electroporations (IUEs) were conducted at embryonic stages E12, E14, and E16 on pregnant mice anesthetized with 3% isoflurane/O₂, maintained at 2%. Following subcutaneous administration of Baytril (5 mg/kg; Bayer) and Meloxicam (300 µg/kg; VITA Laboratories), the abdominal area was sterilized with 70% ethanol and saline. A skin incision exposed the uterine horns, and the lateral ventricle (LV) was visualized using cold light transillumination. A 1-2 µl mixture of StarTrack plasmids (1.8-2 µg/µL in distilled water) was injected into the LV, with Fast Green (0.05%) used to confirm injection accuracy. Electroporation was performed using electrode tweezers (Protech International; #CUY650-P3), applying five square electrical pulses of 50 ms at 950 ms intervals to the dorsal LV, with voltages adjusted for each embryonic stage to minimize developmental risks (E12: 33mV; E14: 35mV; E16: 37mV). After electroporation, embryos were returned to the abdominal cavity, and the incision was closed. Dams were kept in a sterile environment at 37°C post-surgery. Embryos were allowed to develop until analysis at 48 hours post-IUE, P0, or P90.

### 2.4. Episomal copies removal by tamoxifen administration

To activate Cre-recombinase and eliminate non-integrated plasmid copies, tamoxifen (Tx, Sigma-Aldrich T5648-1 G, 20 mg/mL in corn oil) was administered intraperitoneally at 7.5 mg/kg body weight. For embryonic tissues, Tx was given to the pregnant mother 48 hours after IUE. Tx was administered at P8 for adult tissue to avoid complications during parturition.

### 2.5. Tissue processing

For embryonic brain processing, pregnant mice were anesthetized with pentobarbital (Dolethal, 40–50 mg/kg, Vétoquinol) and transcardially perfused with 0.9% saline followed by 4% paraformaldehyde (PFA, Sigma-Aldrich) in 0.1 M phosphate buffer (PB). The uterine horns were then exposed, and embryos were extracted. For E18 embryos, the perfusion was performed with 4% PFA, and brains were post-fixed for 1 hour in the same solution. The brains were then sectioned coronally at 40 or 50 µm using a vibratome and mounted on six gelatinized slides.

For postnatal samples at P90, mice were anesthetized with pentobarbital and transcardially perfused with 4% PFA. The brains were post-fixed for 1 hour with 4% PFA and sectioned coronally at 50 µm using a vibratome with the sections collected in six wells.

### 2.6. Immunohistochemistry

Immunohistochemistry protocols were adapted to the brain’s developmental stage. Embryonic and P0 brain sections were mounted on slides, while postnatal brains were processed with free-floating immunohistochemistry. Sections were first washed and permeabilized in phosphate-buffered saline with 0.5% Triton X-100 (PBS-T), followed by 0.1% PBS-T. Where indicated, sections were treated with 10% methanol for 20 minutes (see Table 1). We performed a heat-induced epitope retrieval using alternating hot and cold citrate buffer (3 x 5 min), when required. Sections were then incubated in blocking buffer (5% normal goat serum, NGS, in 0.1% PBS-T) for at least 2 hours, and primary antibodies diluted in blocking buffer (see Table 1) were applied overnight at 4°C. Following six washes with 0.1% PBS-T, secondary antibodies were added for 2 hours (see Table 1). After washes in 0.1% PBS-T and 1x PBS, sections were mounted on glass slides using Mowiol (Polysciences).

**Table 1.** Primary and secondary antibodies.

| Antibody | Host | Use | Reference | Company |
| --- | --- | --- | --- | --- |
| APC | Ms | 1:300 | OP80 | Calbiochem |
| BLBP | Rb | 1:500 | 32423 | Abcam |
| DCX | Rb | 1:500 | 4604 | Cell signalling |
| GFAP | Rb | 1:1000 | 31745 | Dako |
| KI67 | Rb | 1:300 | AB16667 | Abcam |
| MASH1/ ASCL-1 | Rb | 1:250 | Ab74065 | Abcam |
| NEUN | Ms | 1:500 | MAB377 | Millipore |
| NESTIN | Ms | 1:100 +citrate | 4760 | Cell signalling |
| NG2 | Rb | 1:100 | AB5320 | Millipore |
| OLIG 2 | Rb | 1:500 | AB9610 | Millipore |
| PDGFRA | Rb | 1:300 +MetOH | 31745 | Cell signalling |
| S100B | Rb | 1:500 | AB41548 | Abcam |
| SOX 2 | Rb | 1:200 | 2748 | Cell signalling |
| SOX 10 | Rb | 1:200 | 69661 | Cell signalling |
| TUJ 1 | Ms | 1:300 | MAB1637 | Millipore |
| VIMENTIN | Ms | 1:200 | V6389 | Sigma-Aldrich |
| Alexa Fluor 633 GtRb | Gt | 1:1000 | A-21070 | Thermo Fisher |
| Alexa Fluor 647 GtMs | Gt | 1:1000 | A-21236 | Thermo Fisher |
| Alexa Fluor 647 GtRat | Gt | 1:1000 | A-21247 | Thermo Fisher |
| Alexa Fluor 568 GtRb | Gt | 1:1000 | A-11011 | Thermo Fisher |
| Alexa Fluor 568 GtMs | Gt | 1:1000 | A-11004 | Thermo Fisher |

Ms: mouse, Rb: rabbit, Rt: rat, Gt: goat.

To identify the neural cell types of StarTrack-labeled cells, we used a panel of primary antibodies targeting specific markers for various neural populations (Table 1). Neuroepithelial cells were marked with antibodies against Sox2, Vimentin, and Nestin. Radial glial cells were identified with BLBP and GFAP, while NG2-glia lineage cells were detected with NG2, PDGFRα, and Sox10. Ascl1/Mash1 marked both neuronal and oligodendrocyte precursors. Immature neurons were detected using DCX and TUJ1. The oligodendroglial lineage was labeled with Olig2, while mature oligodendrocytes were identified with APC. Astrocytes were marked with S100β, mature neurons with NeuN, and proliferating cells with Ki67.

### 2.7. Image capture and processing

Sections were examined using epifluorescence microscopy equipped with specific filter cubes for GFP (FF01-473/10), mCherry (FF01-590/20), and Cy5 (FF01-628/40-25). Mosaic images were captured with a confocal microscope (TCS-5 Leica). The excitation (Ex) and emission (Em) wavelengths for each fluorophore were as follows: EGFP (Ex: 488 nm; Em: 498–510 nm, mCherry (Ex: 561 nm; Em: 601–620 nm), and Alexa Fluor 633/647 (Ex: 633 nm; Em: 650–760 nm). Confocal images were processed to generate maximum projection views using LasX software (Leica Application Suite X, Version 3.5.1).

### 2.8. Quantitative analysis and Statistical analysis

Quantitative analyses were conducted using the Cell Counter tool in Fiji (ImageJ, version 1.53q). Sections were placed into six different slides. Focusing on the center of the electroporated area, three representative slices were selected from each slide to analyze embryonic brain samples. In adult mice, the brain was similarly sectioned into six wells, with one section analyzed for every 300 μm of tissue, which included all electroporated slices from one well.

Statistical analyses were conducted using RStudio (version 1.4.1106) and GraphPad Prism (version 6.0, San Diego, CA, USA). Data distribution was assessed for normality with the Kolmogorov–Smirnov test, using the Dallal-Wilkinson-Lillie test for p-values. For non-parametric data, the Mann–Whitney test was employed to compare two groups, while the Kruskal–Wallis test was used for comparisons among more than two groups. Results are presented with a 95% confidence interval, and statistical significance was defined as *p < 0.05, **p < 0.01, ***p < 0.001 and ****p < 0.0001. Data visualization and additional statistical analyses were performed in RStudio to facilitate a comprehensive understanding of the results.

### 2.9. Cell isolation by FAC sorting

At E14, pregnant mice were euthanized by cervical dislocation following approved ethical protocols. The abdominal area was cleaned with 70% ethanol, and the uterine horns were carefully exposed. Embryos were extracted and immediately placed in DPBS (GibcoTM, #14190144). Under a binocular microscope (Nikon, #SMZ1500), the brains were extracted and minced into pieces smaller than 1 mm in FACs medium (HBSS, HEPES, EDTA, BSA, DNAse). To account for biological variability, each sample contained the pooled brains of all embryos from two pregnant mice. The tissue was mechanically dissociated using heat-shrink pipettes, followed by enzymatic digestion with TrypLE Express (GibcoTM, #12604013) for 2 minutes at 37°C. After enzyme inactivation, the cell suspension was resuspended in FACS buffer and treated with red blood lysis buffer (Myltenyi, #130-094-183). To preserve RNA integrity, RNAse inhibitor (Applied Biosystems™, #N8080119) was added to the cell suspension. The mixture was then filtered through a 40 µm pluri-strainer (pluriSelect, #43-10040-40) before being sorted using a FACSAria Flow Cytometer (BD) at the CBM (Madrid, Spain). EGFP+ cells were isolated and collected into a Lysis Buffer for RNA isolation. The working area and equipment were cleaned with RNAseZap (Nzytech, #MB16001) and 70% ethanol to prevent RNA degradation or contamination.

### 2.10. RNA isolation

RNA isolation was performed using the RNAqueous™ Micro Total RNA Isolation Kit (Invitrogen™, #AM1931). The cells were lysed with lysis buffer and vortexed, followed by the precipitation of RNA adding 100% ethanol. The resulting lysate was loaded onto a microfilter cartridge, washed, and treated with DNase using the PureLink™ DNase Set (Invitrogen™, #12185010) to eliminate residual DNA. After an additional wash, RNA was eluted with nuclease-free water. All surfaces and materials were treated with RNAseZap (Nzytech, #MB16001) and ethanol to ensure the integrity of the RNA. The concentration and integrity of the RNA samples were assessed using a Bioanalyzer (Agilent, 4200 TapeStation System).

### 2.11. Library preparation and RNA sequencing

RNA samples were amplified using SMARTer amplification (Novogene, UK). After quality assessment, mRNA libraries were prepared with Poly A enrichment and sequenced on the NovaSeq™ System PE150 (Novogene, UK), yielding 6 GB of raw data per sample. A total of ten samples were analyzed, including five from NG2-NPCs and five from RGPs.

### 2.12. Bioinformatic analysis of RNA-Seq data

The RNA sequencing data were processed and analyzed using a standardized pipeline. First, raw sequence reads in fastq format underwent initial quality control through in-house Perl scripts. Clean reads were obtained by removing reads containing adapters, poly-N sequences, and low-quality. To ensure data quality Q20, Q30, and GC content were calculated by Novogene. All downstream analyses were based on high-quality clean reads. Cleaned reads were aligned to the reference genome (Mus musculus, GRCm38.p6) using HISAT2 (version v2.0.5). Aligned reads were quantified at the gene level using featureCounts (version 1.5.0-p3). Expression levels were calculated as FPKM (Fragments Per Kilobase of transcript per Million mapped reads), considering both sequencing depth and gene length.

After initial quality filtering and preprocessing of the RNA-seq data, we conducted dimensionality reduction using t-distributed Stochastic Neighbor Embedding (t-SNE) to visualize the overall distribution and relationships among gene expression profiles. Subsequently, we employed the (Hierarchical Density-Based Spatial Clustering of Applications with Noise (HDBSCAN) algorithm for clustering the data, to identify distinct groups of samples based on their expression patterns. According to this analysis, two samples were considered outliers as they did not cluster with the corresponding group.

For differential gene expression analysis, gene count matrices were analyzed with DESeq2 (version 1.44.0) in the R environment (39). Genes with low counts (fewer than 10 reads across all samples) were filtered out. Normalization was performed using the median of ratios method. Differential expression analysis was conducted comparing NG2+cells vs GFAP+cells. Results were reported as log2 fold changes (log2FC) with associated p-values and adjusted p-values using the Benjamini-Hochberg method to control for the false discovery rate (FDR). Genes with a p-value < 0.05 were considered significant.

Gene Ontology (GO) analyses were conducted using clusterProfiler (version 4.12.6)(40), with a p-value cut-off of 0.05 and a q-value cut-off of 0.05 to identify significantly enriched biological processes. Additionally, the simMatrix function was used to calculate the semantic similarity between GO terms (“Wang” method and threshold = 0.80). GO terms visualization and relationships were carried out using the GOplot R package (version 1.0.2).

KEGG pathway enrichment analysis was performed using ClusterProfiler in R. Differentially expressed genes (DEGs) were mapped to KEGG pathway IDs, and significant pathways were identified with an adjusted p-value < 0.05. Pre-ranked gene set enrichment analysis (GSEA) was used to interpret gene expression data in terms of biological processes (41). All statistical analyses and visualizations were conducted in R (version 2024.04.2).

## 3. Results

### 3.1. Temporal Patterning Reveals Distinct Spatial Localization of NG2+cells compared to GFAP+cells

Our study investigates the differences between NG2-progenitors (NG2-NPCs) and GFAP+ progenitors (GFAP-NPCs/ RGCs) during embryonic development, focusing on their cellular, molecular, and functional characteristics. Specifically, we sought to determine whether these progenitors represent distinct populations and how these differences might influence cell fate decisions during development. To this end, we analyzed the presence and characteristics of NG2-NPCs and RGCs across embryonic stages. Using co-electroporation of plasmids driven by GFAP or NG2 promoters, we examined the co-localization of NG2+cells and GFAP+cells at E12, E14, and E16 (Figure 1A). Control experiments for co-electroporation were performed to validate the approach (Figure S1). To minimize biases related to plasmid integration, we employed two complementary experimental strategies: GFAP-GFP/ NG2-mCherry-StarTrack (Figure 1A, left) and GFAP-mCherry/ NG2-EGFP-StarTrack (Figure 1A, right). Labeled cells were quantified 48 hours post-*in utero* electroporations (IUE).

**Figure 1.**
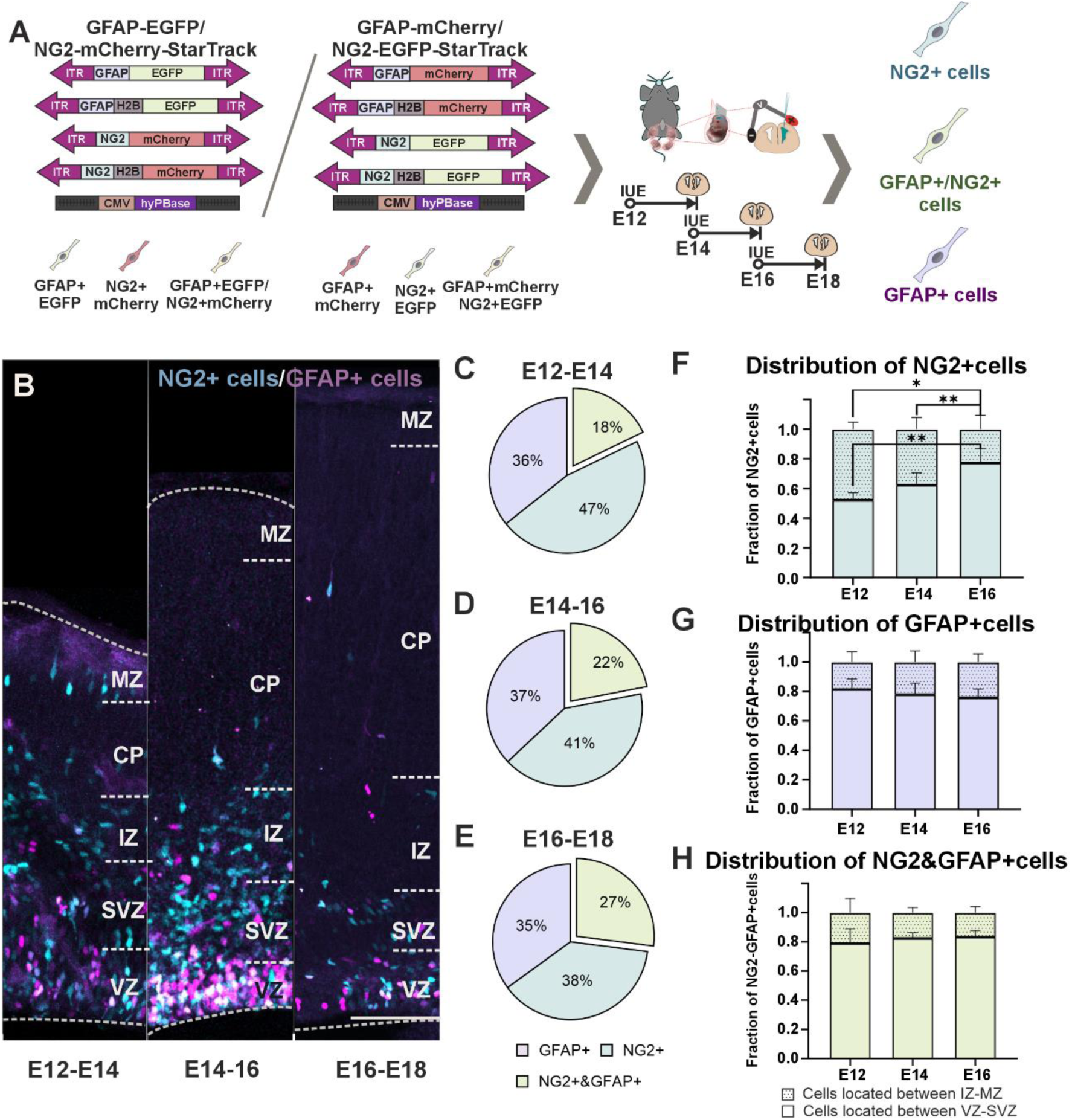
Temporal Patterning of NG2+cells vs. GFAP+cells during brain development. (A) Co-labeling of NG2+cells and GFAP+cells after *in utero* electroporation of StarTrack plasmids driven by the NG2 or GFAP promoter. (B) Representative images of NG2+cells (cyan) and GFAP+cells (magenta) at various developmental stages. (C-E) Fraction of NG2+cells (blue), GFAP+cells (purple), and double-positive (NG2+&GFAP+) cells (green) among all StarTrack-labeled cells analyzed 48 hours post-targeting progenitors at E12, E14, and E16. (F-H) Spatial distribution pattern of NG2+cells (F), GFAP+cells (G), and NG2&GFAP+cells (H) within the ventricular zone (VZ) and subventricular zone (SVZ), as well as between the intermediate zone (IZ) and the marginal zone (MZ). N = 6 mice per experimental condition. * p<0.05, ** p < 0.01. Scale bar: 100 μm.

Our findings revealed that NG2+ and GFAP+ cells largely represented distinct progenitor populations but a smaller subset co-expressed both markers during the development (Figure 1B). Quantitative data indicated that the proportion of double-positive (NG2+&GFAP+) cells was relatively low, comprising 18% of the population at E12, 22% at E14, and 27% at E16 (Figures 1C-E). However, most cells expressed NG2 (38-47%) or GFAP (35-37%) across all stages. Notably, the fraction of NG2+cells was significantly larger than NG2+&GFAP+cells after targeting NPCs at E12 (p= 0.0002) and E14 (p=0.0283, *data not shown*). In contrast, no differences were found between the fraction of GFAP+cells and NG2+&GFAP+cells (p= 0.116 at E12 and p=0.1547 at E14, *data not shown*).

The spatial distribution of these populations also differed depending on the type of NPCs and developmental stage (Figures 1F-H). Forty-eight hours after targeting, GFAP+cells (Figure 1G) and NG2+/GFAP+cells (Figure 1H) were predominantly located near the lateral ventricle (LV), with a 70-80% of cells remaining close to the LV across all stages (p= 0.0006 at E12, p=0.0022 at E14 and E16). In contrast, NG2+cells exhibited a distinct distribution pattern at earlier stages (Figure 1F). After targeting NPCs at E12, NG2+cells were evenly dispersed between the ventricular-subventricular zone (VZ-SVZ) and the intermediate and marginal zones (IZ-MZ) with no significant differences between these two regions (p=0.0728). However, by E14, NG2+cells began to concentrate near the LV (60%, p=0.0022), a pattern that persisted and aligned more closely with GFAP+cells and double-positive cells by E16 (80%, p=0.0022) (Figure 1F).

These results highlight significant differences in the presence and spatial localization of NG2+cells and GFAP+cells during early embryonic development. The distinct temporal and spatial distribution patterns suggest the existence of unique progenitor populations, each with potentially different roles in central nervous system development.

### 3.2. Functional Divergence in Long-Term Differentiation of NG2 vs. GFAP Progenitors

To determine whether the initial differences in cellular populations result in functional distinctions, we analyzed the differentiation capacities of NG2-NPCs and GFAP-NPCs at various embryonic stages, specifically focusing on short-term (P0) and long-term outcomes (P90) (Figure 2A). To specifically label NG2-NPCs or GFAP-NPCs, we performed IUE at E12, E14, and E16 using UbC-(NG2 or GFAP-PB)-EGFP-StarTrack which include specific transposases (NG2-hyPBase or GFAP-hyPBase), along with UbC-EGFP and UbC-H2B-EGFP-StarTrack plasmids (12,13,33). This strategy allowed us to trace the progeny of NG2-NPCs or GFAP-NPCs over time. In the progeny of both types of progenitors, we identified several cell types, including neuronal cells (Figure 2Ba) as well as glial lineages, such as astrocytes (Figure 2Bb), NG2-glia (Figure 2Bc), and oligodendrocytes (Figure 2Bd).

**Figure 2.**
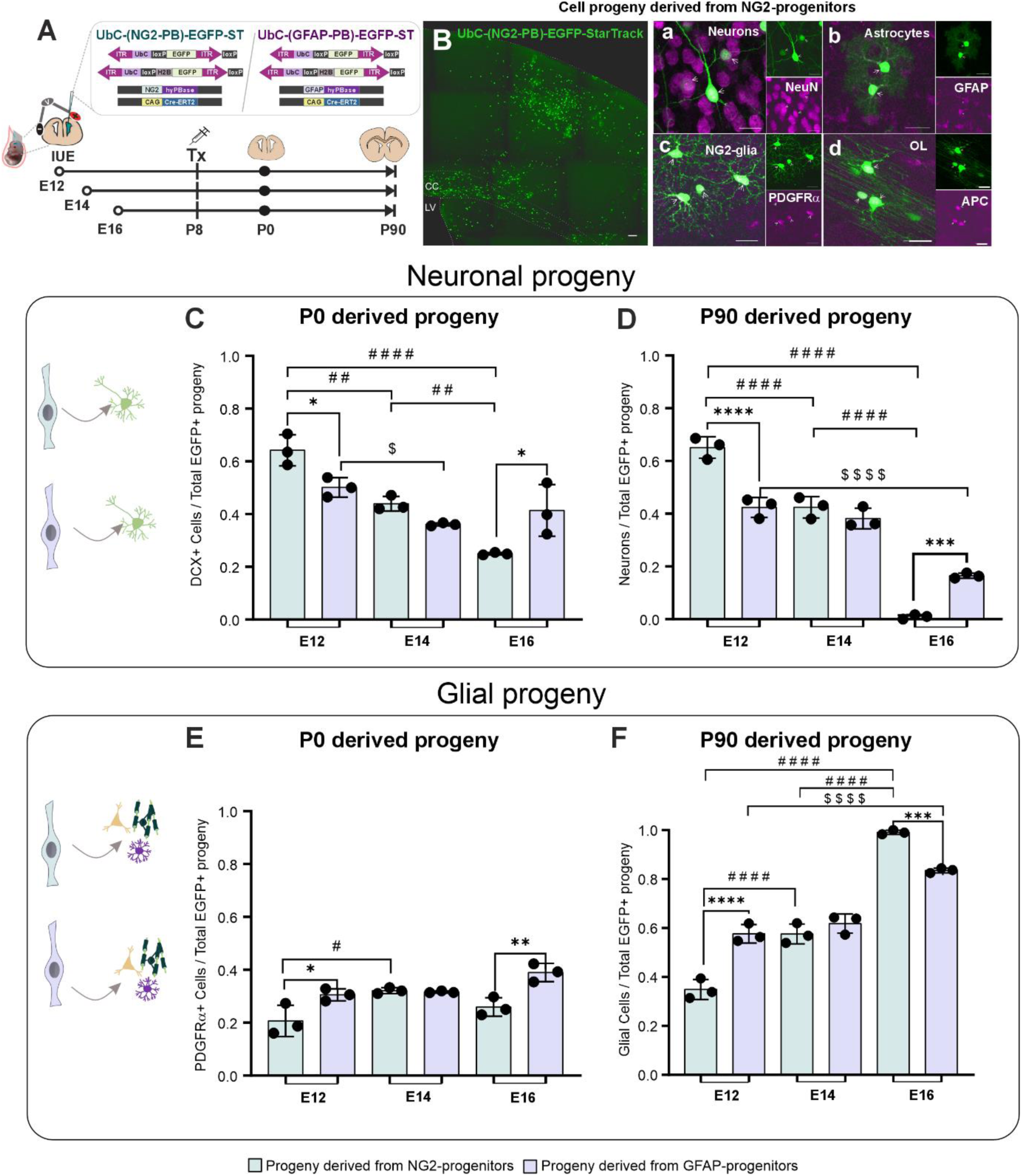
Cell progeny of Embryonic NG2-NPCs vs. GFAP-NPCs. (A) Experimental design for *in utero* electroporation at E12, E14, and E16 using UbC-(NG2 or GFAP-PB)-EGFP-StarTrack plasmid to tag NG2-NPCs or GFAP-NPCs, respectively, and their complete cell progeny at short (P0) and long-term (P90). (B) Representative images of cell types derived from NG2-NPCs, with magnified images of NeuN+ neurons (a), S100β+ astrocytes (b), PDGFRα+ NG2-glia (c) and APC+ oligodendrocytes (OL) (d). (C-D) Quantification of neuronal cells in all EGFP+cells derived from NG2-NPCs (blue) and GFAP-NPCs (purple) at P0-DCX+ (C) and P90 (D). (E-F) Fraction of glial cells in all EGFP+ cells derived from NG2-NPCs (blue) and GFAP-NPCs (purple) at P0 (E) and P90 (F). N = 3 mice per experimental approach. Asterisks (*) indicate significant differences comparing NG2 vs GFAP-derived progeny, hashtags (#) differences in NG2-NPC progeny at different stages, and dollar signs ($) are used to indicate significant differences in GFAP-NPC progeny along the development. * p<0.05, ** p < 0.01. *** p <0.001, **** p < 0.0001 Scale bar in B: 100 μm; Ba-d: 25μm.

#### 3.2.1. Neurogenic potential

Our analysis revealed that NG2-NPC progeny has a significantly higher fraction of neuronal cells when labeled at both P0 (64,2%) and P90 (65,1%) when targeted at E12. However, the neuronal output decline significantly at later stages, particularly at E16 at both short (24%) and long-term (0,9%) (E12 vs E14: p = 0.0038 at P0, p <0.0001 at P90; E12 vs E16: p<0.0001 at P0 and P90) (Figures 2C and D). A similar decline was also observed in GFAP-NPC P90 progeny but only between E12 and E16 (p < 0.0001) (Figures 2C and D). The difference in neuronal output between both types of NPCs was most evident at E12, where NG2-NPCs produced 65,1% of neuronal cells compared to 42.4% for GFAP-NPCs (p = 0.0399 at P0; p < 0.0001 at P90) (Figures 2C and D). By contrast, targeting NPCs at E16, GFAP-NPCs exhibited a higher neuronal output than NG2-NPCs at both P0 (41,4% vs. 24%, p = 0.0162) and P90 (16,5% vs. 0,9%, p = 0.0003) (Figures 2C and D). These findings indicate that NG2-NPCs possess greater neurogenic potential during early developmental stages, while GFAP-NPCs maintain a more prolonged differentiation pattern, with a delayed decline in neurogenic potential.

#### 3.2.2. Gliogenic potential

Regarding gliogenesis, the GFAP-NPCs progeny at P0 displayed a significantly higher percentage of PDGFRα+ cells compared to NG2-NPCs at E12 (30,5% vs. 20,7%, p = 0.035) and E16 (39,0% vs. 25,9%, p = 0.0056) (Figure 2E). This trend persisted at P90, where GFAP-NPCs labeled at E12 produced significantly more PDGFRα+ cells than NG2-NPCs (GFAP-NPCs: 57,6% vs. NG2-NPCs: 34,9%, p < 0.0001) (Figure 2F). Interestingly, at E16, the glial output of NG2-NPC progeny at P90 surpassed that of GFAP-NPCs (97,0% vs. 81,1%, p = 0.0003) (Figure 2F).

These results reveal distinct temporal differentiation patterns between NG2-NPCs and GFAP-NPCs, underscoring their specialized roles in neurogenesis and gliogenesis throughout central nervous system development. NG2-NPCs demonstrate stronger early neurogenic potential with a progressive decline and a shift towards gliogenesis at later stages. In contrast, GFAP-NPCs maintain a more prolonged neurogenic phase, with a delayed transition to gliogenesis. This dynamic interplay between NG2-NPCs and GFAP-NPCs suggests complementary roles in balancing neurogenesis and gliogenesis during CNS development.

### 3.3. Distinct Molecular Signatures of NG2 and GFAP Progenitors

These progenitors were detailly characterized to elucidate the molecular basis underlying the functional divergent roles between NG2-NPCs and GFAP-NPCs during early brain development. After targeting NG2-NPCs and GFAP-NPCs through the combined use of transposase systems and StarTrack plasmids, driven by specific promoters tailored for each progenitor type, we characterized them using both immunohistochemistry and transcriptomic approaches (Figures 3A-B).

**Figure 3.**
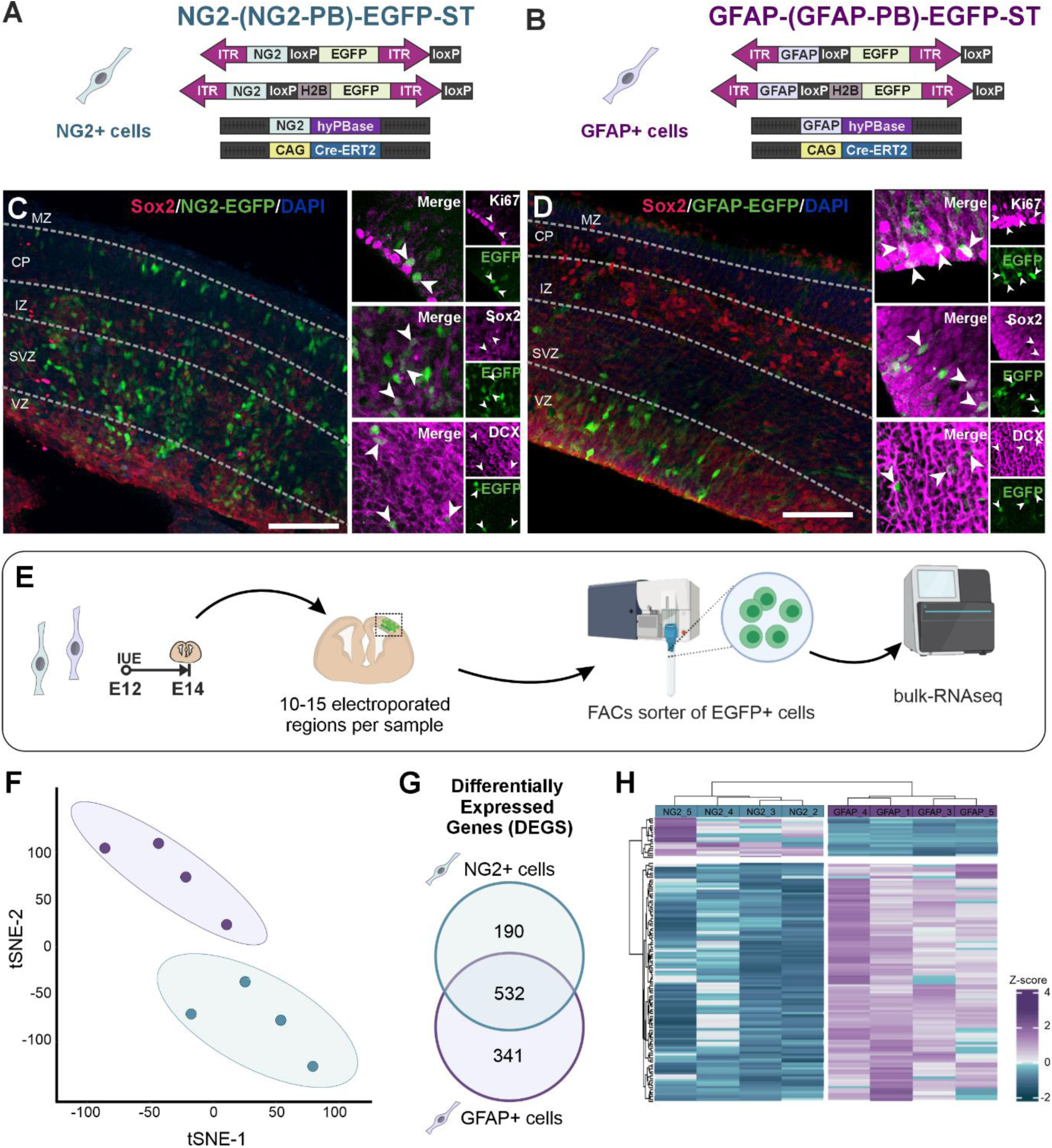
Transcriptomic characterization of NG2+cells and GFAP+cells at earlier stages. (A-B) Schematic representation of StarTrack plasmids used to label NG2+cells (A) and GFAP+cells (B). (C-D) Immunohistochemical staining for proliferation (Ki67) and progenitor markers (Sox2), and neuroblast (DCX) after targeting NG2+cells (C) and GFAP+cells (D). (E) Experimental workflow for bulk-RNA sequencing of E14 cells isolated by FACS sorting after targeting them at E12 by IUE. (F) t-SNE plot of NG2+cells (blue) and GFAP+cells (purple) based on their transcriptomic profiles. (G) Venn diagram representing differentially expressed genes (DEGs) between NG2+cells and GFAP+cells. (H) Heatmap of DEGs between NG2+cells and GFAP+cells. Scale bar: 100 μm.

While we did not notice clear differences between NG2+ and GFAP+ populations in common progenitor markers expression using immunohistochemical staining, such as Sox2 and Nestin (Figures 3C-D), the transcriptomic analysis provided deeper insights into their molecular divergence. To capture the early transcriptional profiles of these progenitors during a critical developmental window, before their divergence into distinct lineages, we performed IUE at E12 to label NG2+cells and GFAP+cells (Figure 3E). Forty-eight hours later, the labeled cells were isolated via fluorescence-activated cell sorting (FACS), followed by bulk-RNA sequencing to generate a detailed transcriptomic profile of each cell population (Figure 3E). This integrative approach enabled us to isolate NG2+cells and GFAP+cells, providing a deeper understanding of the intrinsic molecular differences that could explain the distinct functional roles and behaviors observed in these progenitor populations.

The transcriptomic analysis revealed both similarities and distinct molecular signatures that differentiate NG2+ cells from GFAP+ cells. t-SNE and HDBSCAN clustering analysis separated and grouped the two populations, highlighting significant differences in their gene expression profiles (Figure 3F). A total of 1,063 differentially expressed genes (DEGs) between NG2+ cells and GFAP+ cells were statistically significant. Of these, 190 genes were uniquely enriched in NG2+ progenitors, while 341 genes were uniquely enriched in GFAP+ progenitors (Figure 3G). Heatmap representation based on the z-scores of these DEGs highlighted the distinct transcriptional programs for each population (Figure 3H). These findings underscore that, despite the similar expression of common progenitor markers observed through immunohistochemistry, NG2+ and GFAP+ progenitors are also molecularly distinct by E14. This molecular divergence may underlie their divergent roles and behaviors during central nervous system development.

Deepening the analyses of DEGs in NG2+cells compared to GFAP+cells, the volcano plot displayed genes that were significantly upregulated (adjusted p ≤ 0.05, logarithm fold change (log2FC) ≥ 1) or downregulated (adjusted p ≤ 0.05, log2FC ≤ −1) in NG2+cells (Figure 4A). Specifically, 341 genes had a logFC < −0.5 (a 0.67-fold reduction), with 109 of these showing their expression reduced by half (logFC < −1) in NG2+cells compared to GFAP+cells (Figure 4B). Conversely, 190 genes exhibited a logFC > 0.5 (1.4-fold increase), with 19 of these presenting a doubling of expression (logFC > 1) (Figure 4B). These findings unveil distinct gene expression profiles between the two cell populations, suggesting underlying regulatory differences. A substantial proportion of the DEGs encoded proteins (45%), while 10% were non-coding genes. The rest of the DEGs (45%) were uncharacterized (Figure 4C). Most upregulated genes in NG2+cells were protein-coding (Figure 4D), while downregulated genes reveal more variability, including non-coding RNA, protein-coding genes, and a great percentage of unknown genes (Figure 4E). Among these uncharacterized genes, we identified predicted gene models (*Gm00000*), indicating the potential presence of novel, unidentified factors that may play crucial roles in the distinct behaviors and lineage choices of NG2-NPCs and GFAP-NPCs.

**Figure 4.**
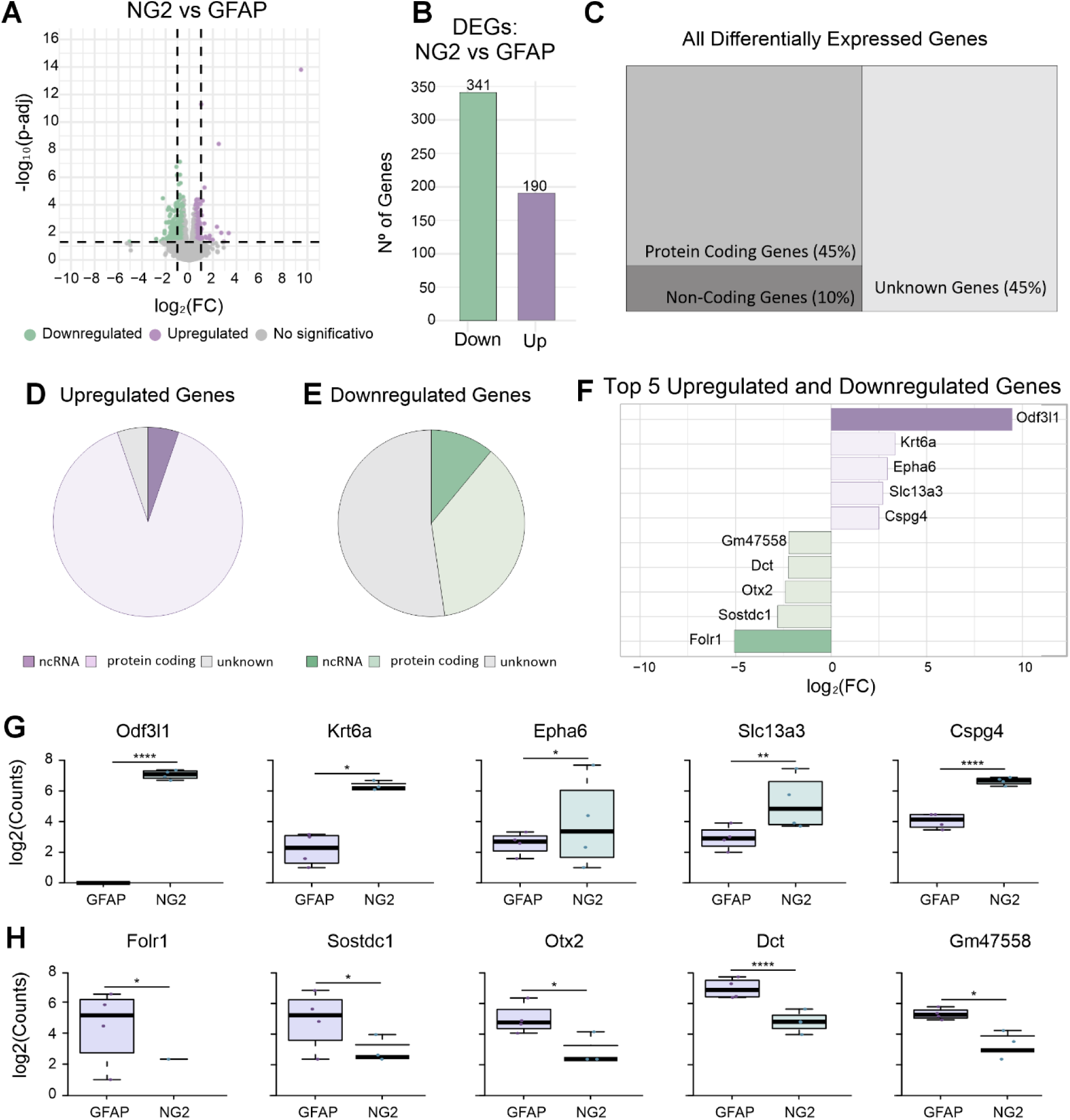
Differential gene expression analysis between NG2+cells and GFAP+cells. (A) Volcano plot displaying differentially expressed genes (DEGs) between NG2+cells and GFAP+cells. Green dots represent downregulated genes (logFC < −1), purple dots represent upregulated genes (logFC > 1), and grey dots represent non-significant genes. (B) Bar plot of the total number of upregulated and downregulated genes comparing NG2+cells vs GFAP+cells, considering a logFC > 0.5 or <-0.5, respectively. (C-E) The proportion of protein-coding, non-coding, and unknown function genes among all differentially expressed genes (C), upregulated genes (D), and downregulated genes (E). (F) Top five upregulated (right-purple) and downregulated (left-green) genes in NG2+cells compared to GFAP+cells. (G-H) Counts (Log2) of top upregulated (G) and downregulated (H) genes in NG2+cells (blue) compared to GFAP+cells (purple). * p<0.05, ** p < 0.01. *** p <0.001, **** p < 0.0001.

The most upregulated genes in NG2+cells were *Odf3l1*, *Krt6a*, *Epha6*, *Slc13a3*, and *Cspg4* (Figures 4F and 4G). *Odf3l1* (*Cimap1c*) encodes a ciliary microtubule-associated protein involved in cell remodeling, implying a role in cytoskeletal dynamics during cell differentiation (Figure 4G). *Krt6a* (Keratin 6a), a member of the keratin family, is transiently expressed during neuronal differentiation, highlighting its potential importance in neurogenesis and in the structural reorganization of cells transitioning from progenitors to mature neurons (Figure 4G). *Epha6* (Eph receptor A6), part of the ephrin receptor family, is expressed in many neuronal subtypes, playing a critical role in axonal guidance and synaptic connectivity (Figure 4G). *Slc13a3*, a sodium-dependent dicarboxylate transporter, is essential for transporting key neuronal metabolites, implicating its involvement in neuronal metabolic support (Figure 4G). Finally, *Cspg4* encodes the proteoglycan Nerve/Glia 2 (NG2), a marker for NG2-glia or oligodendrocyte precursor cells (OPCs), emphasizing their role in myelination and brain plasticity (Figure 4G).

Conversely, the most downregulated genes in NG2+cells included *Folr1*, *Sostdc1*, *Otx2*, *Dct*, and *Gm47558* (Figures 4F and 4H). *Folr1* encodes folate receptor alpha, which plays a crucial role in cell proliferation and folate metabolism, essential for neurodevelopment (Figure 4H). *Sostdc1*, a sclerostin domain protein, is an antagonist of bone morphogenetic proteins (BMPs) that regulate neural progenitor cell proliferation (Figure 4H). This downregulation suggests reduced BMP signaling, which may be linked to differentiation processes in NG2+ cells. *Otx2* (orthodenticle homeobox 2) is involved in the early neuroectoderm specification and becomes downregulated in the dorsal telencephalon, indicating a transition away from early neurodevelopmental stages (Figure 4H). Lastly, *Dct*, encoding dopachrome tautomerase, is implicated in neural stem cell proliferation, supporting the notion that NG2+ cells have a reduced proliferative capacity compared to GFAP+ cells (Figure 4H).

Gene Ontology (GO) enrichment analysis highlights key significant biological processes that are either increased or decreased in NG2+cells compared to GFAP+cells. Processes such as *’developmental maturation (GO:0021700)’*, *’vesicle-mediated transport in synapses (GO:0099003)’, ‘cognition GO:0050890* and *‘regulation of postsynaptic membrane neurotransmitter receptor levels (GO:0099072)’* were significantly increased in NG2+cells (Figures 5A and B). These results align with the higher expression of genes involved in neurogenesis and synaptic formation, suggesting that embryonic NG2+cells are more actively engaged in shaping early neuronal connections and playing an essential role in establishing the brain’s wiring. Conversely, processes such as ‘*forebrain development (GO:0030900)’*, *’neural precursor cell proliferation (GO:0061351)’*, and *’maintenance of cell number (GO:0098727)’* were significantly decreased in NG2+cells. This indicates that NG2+cells are less committed to maintaining progenitor stages and are more inclined towards differentiation.

**Figure 5.**
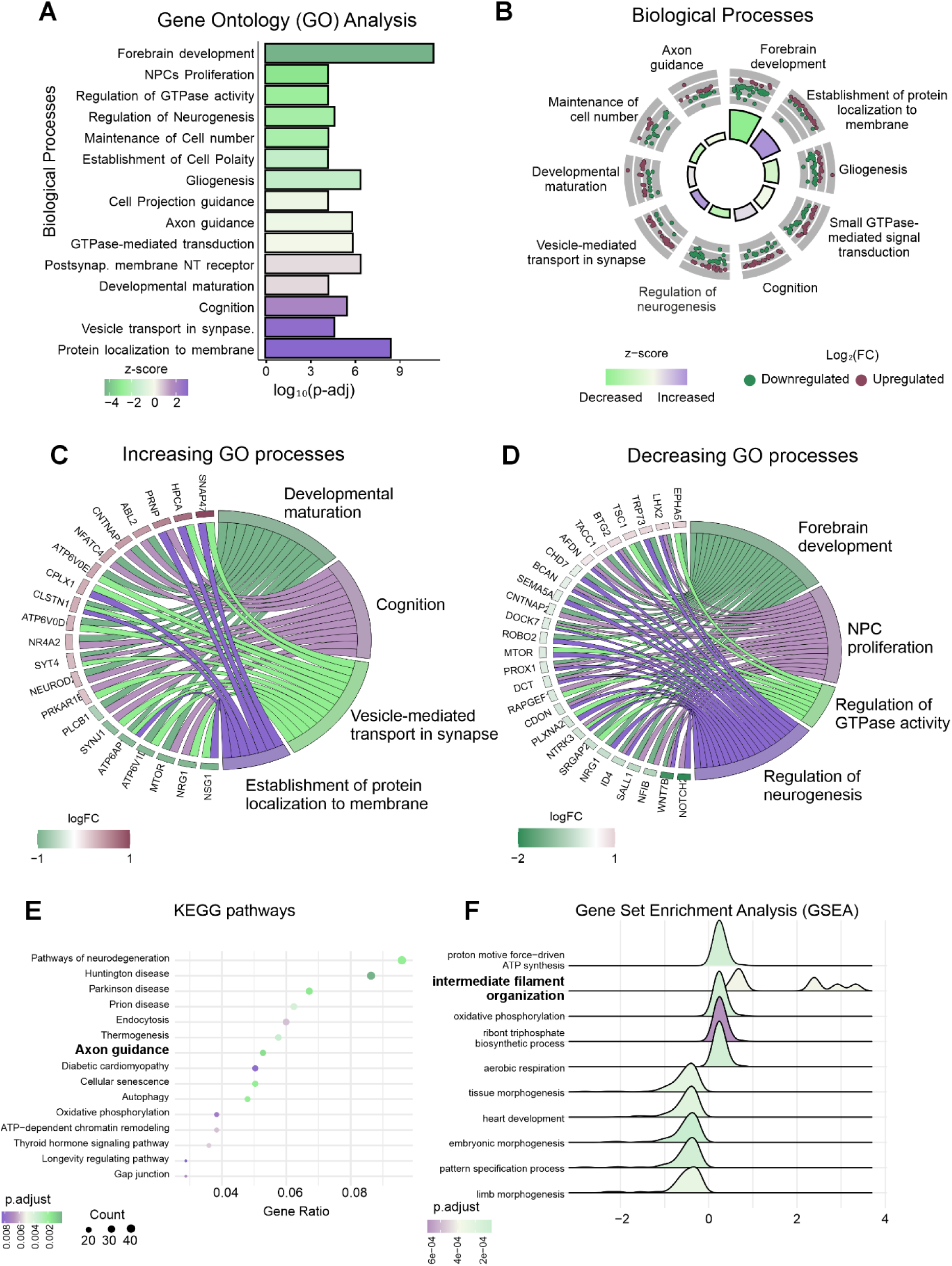
Gene Ontology (GO) and KEGG pathway analysis of differentially expressed genes between NG2+cells and GFAP+cells. (A) GO biological process downregulated (light green) and upregulated (purple) in NG2+cells compared to GFAP+cells. (B) A circular GO plot of key biological processes, highlighting increased and decreased processes in NG2+cells relative to GFAP+cells, including upregulated (red points) and downregulated (green points) genes. (C, D) Chord diagrams illustrate the genes contributing to the increasing (C) and decreasing (D) GO processes. (E) KEGG pathway enrichment in NG2+cells compared to GFAP+cells. (F) Gene Set Enrichment (GSEA) analysis in NG2+cells compared to GFAP+cells.

To explore further the increased processes in the NG2+ cell population, we focused on those with the highest Z-scores (Figure 5C), analyzing genes involved in at least two of these processes. Notably, we observed genes such as *Calsyntenin 1 (Clstn1)*, implicated in *’developmental maturation’*, *’vesicle-mediated transport in synapses’*, and *’establishment of protein localization to membranes’*. *Clstn1* is known to play a role in axon guidance and is involved in neural circuit formation before the onset of synaptogenesis. We also identified *ATPase, H+ transporting V0 subunit D1 (Atp6v0d1)*, which in neural precursors of the developing mouse cortex has been shown to deplete neural stem cells by promoting their differentiation and the generation of neurons. Another key gene, *Snap47 (synaptosomal-associated protein, 47)*, is implicated in neuronal morphogenesis. Furthermore, *NeuroD2* is involved in developmental maturation and cognition, contributing cell remodeling processes.

In contrast, when examining processes with lower Z-scores (Figure 5D), we found genes such as *LIM Homeobox 2 (Lhx2)*, *BTG Anti-Proliferation Factor 2* (*Btg2)*, *Brevicam* (*Bcan)*, *Prospero Homeobox 1 (Prox1)*, *Cdon*, *Inhibitor of DNA Binding 4 (Id4)*, and *Notch2* associated with ‘*forebrain development’*, *‘neural precursor cell proliferatio’*, and *‘regulation of neurogenesi’*. *Lhx2* regulates cortical progenitor proliferation by regulating the expression of *Hes1*, a *Notch* signaling pathway effector. *Btg2, Cdon* and *Notch2* are also associated with cell proliferation and maintenance of RGCs. This pattern reinforces the divergence in developmental roles between NG2+cells and GFAP+cells, suggesting that these genes may contribute to the progenitor identity and maintenance functions more characteristic of GFAP+cells.

The KEGG pathway enrichment analysis (Figure 5E) further revealed specific pathways differentially regulated in NG2+cells. Among the significantly upregulated pathways in NG2+cells, we found “*axon guidance (mmu04360)”* and “*ATP-dependent chromatin remodeling (mmu03082)”*. Enriching “*axon guidance”* pathways strengthens the hypothesis that NG2+cells are essential in establishing neural connectivity during this critical development period. The involvement of “*chromatin remodeling”* pathways indicates that NG2+cells may also contribute to cellular robustness and plasticity, potentially enhancing their ability to adapt and support neurogenesis. Complementing this, Gene Set Enrichment Analysis (GSEA) (Figure 5F) identifies enriched gene sets like “*ATP synthesis*”, “*intermediate filament organization*”, and “*tissue morphogenesis*”, pointing to increased metabolic activity, cytoskeletal reorganization, and developmental patterning in NG2+cells compared to GFAP+cells.

Together, these analyses highlight the molecular and functional differences that may influence the distinct developmental and proliferative behaviors of NG2+ and GFAP+ progenitor populations. The GO and KEGG pathway analyses provide valuable insights into the specialized functions of these progenitors in brain development, revealing how each cell type uniquely contributes to neural circuit formation and the maintenance of cellular proliferation. During early development, NG2+cells exhibit transcriptional profiles that favor neurogenesis and synaptic development, indicating their involvement in constructing neural circuits. In contrast, GFAP+cells maintain gene expression profiles that sustain progenitor identity and promote proliferation. These insights deepen our understanding of the mechanisms by which these progenitor populations contribute to brain development and lineage specification, underscoring their unique roles in neural architecture and cellular diversity.

## 4. Discussion

NG2+cells have traditionally been classified as oligodendrocyte precursors. However, recent studies, including ours, support that these cells have broader neurodevelopmental potentials. In this project, we focused on unraveling the functional roles of NG2-expressing progenitors (NG2+cells) and their relationship with radial glial cells (RGCs, GFAP+cells) during embryonic development. Combining genetic labeling techniques and transcriptomic analysis, we explored the spatial, molecular, and functional differences between these two progenitor populations. Our findings reveal distinct developmental trajectories, cellular outputs, and molecular signatures, shedding light on their complementary roles in CNS development.

### 4.1. Divergence Developmental Trajectories

Our data not only demonstrate that the two progenitor populations (NG2+ and GFAP+) occupy distinct spatial niches in the developing cortex, but also follow divergent differentiation trajectories, starting as early as E12 (Figure 6). This spatial segregation likely reflects the unique developmental and proliferative stages of each progenitor type and specialized functional roles in cortical assembly. Initially, NG2+cells are broadly distributed across cortical regions, while GFAP+cells are localized near the ventricular zone (VZ), adjacent to the lateral ventricles. As development progresses, NG2+cells gradually converge spatially with GFAP+cells, with both populations gathering within the VZ. This spatial convergence could indicate a shift in the functional roles of these cells, aligning them together in the VZ to better respond to region-specific cues crucial for late-stage cortical maturation.

**Figure 6.**
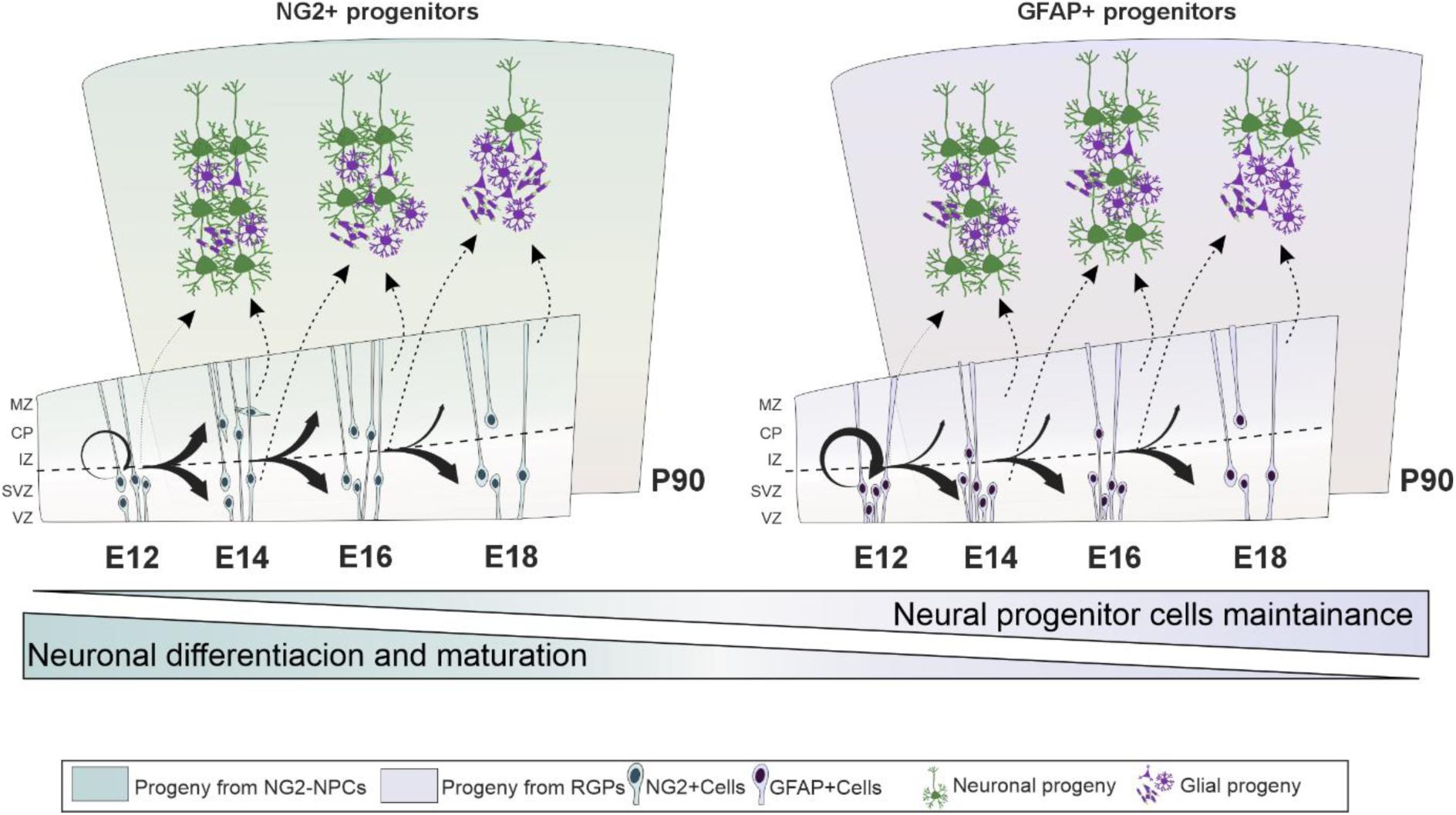
Temporal dynamics of neuronal and glial lineage differentiation from NG2-NPCs and GFAP-NPCs. Differentiation trajectories of NG2-NPCs (left panel) and GFAP-NPCs (right panel) from embryonic stages (E12, E14, E16) to adulthood (P90). Both progenitor populations contribute to neuronal and glial cell lineages, but their differentiation potentials change over time. NG2-NPCs exhibit a prominent shift from neurogenesis to gliogenesis, with increased expression of neurogenesis-related genes at early stages. In contrast, GFAP-NPCs display a delayed transition to gliogenesis, maintaining a more progenitor-like transcriptomic profile at earlier developmental stages.

These spatially distinct niches may also influence the timing and nature of fate decisions within each progenitor population. Our results align with prior studies showing that apical radial glial cells (aRGCs) in the VZ act as primary NSCs, characterized by apico-basal polarity and interkinetic nuclear migration. These cells primarily give rise to basal progenitors (BPs) outside the VZ (42–44). The proximity of GFAP+cells to the VZ suggests they are more directly influenced by signals from this niche, which may support their progenitor potential and contribute to neurogenesis. As development continues, these GFAP+cells transition to produce glial cells. In contrast, NG2+cells, which are initially more dispersed throughout the cortex, could respond, to cues from the intermediate zone or cortical plate, producing cell types that align with the developmental needs of each stage. Their commitment to VZ later in development could represent a shift in function as neurogenesis declines and gliogenesis becomes more prominent.

The presence of a subset of cells (∼20%) expressing both NG2 and GFAP markers across all developmental stages could indicate the existence of an intermediate or transitional cell state within the progenitor pool, enhancing the adaptability of the cortex. This dual-marker population may act as a flexible reservoir, bridging the neurogenic and gliogenic phases or maintaining the progenitor pool while adapting to cortical needs and the contextual cues they receive.

The observed heterogeneity in cortical precursor cells supports various models of progenitor behavior (45–51). One model proposes that NSCs undergo temporal shifts in fate, initially producing neurons for the successive cortical layers and later transitioning to a gliogenic phase (50,51). An alternative hypothesis argues that NSCs represent a multipotent pool, with each progenitor responding to intrinsic and extrinsic cues to generate different neuronal or glial subtypes, sequentially recruiting distinct progenitors as development progresses (46,48). In this context, our findings suggest that NG2+ progenitors may possess a high degree of flexibility to respond to signals from different cortical regions, adjusting their developmental output accordingly. In contrast, GFAP+ progenitors may serve as a more stable, consistent source of neural cells, maintaining structural integrity and ensuring continuity within the progenitor pool over time.

Our study highlights the functional divergence and complementary roles of NG2+ and GFAP+ progenitors in CNS development. By understanding those progenitorś trajectories and molecular signatures, we can gain deeper insights into the complex processes that govern neurogenesis and gliogenesis in the developing brain.

### 4.2. Temporal Dynamics of NG2 and GFAP Progenitors: Divergent and Complementary Roles in Neurogenesis and Gliogenesis

NG2-NPCs and GFAP-NPCs contribute to the generation of neurons and glial cells, in alignment with the general pattern of cortical development, where neurogenesis predominates at earlier stages, followed by overlapped gliogenesis later (7,17,52). However, our findings reveal significant differences in their temporal patterns and differentiation potentials, highlighting their complementary roles in CNS development.

NG2-NPCs exhibit robust neurogenic potential when labeled early (E12), producing a substantial percentage of neuronal progeny at P90, with neuronal fate already determined at P0. However, their neurogenic output significantly declines at later stages marking a shift towards gliogenesis. This temporal shift aligns with the “common progenitor model” (51,53) implying their dual role in early circuit formation and subsequent glial network development. This fate versatility of NG2+cells has been observed in adult environments where NG2+cells generate oligodendrocytes but also astrocytes and neurons (26,27,29–32). In contrast, GFAP-NPCs, often associated with radial glial cells (RGCs), maintain a more stable neurogenic phase, with a delayed decline in neuronal output from E12 to E16. GFAP-NPCs compared to NG2-NPCs, show a prolonged neurogenic capacity but with a lower yield of neurons at E12 and maintaining neurogenic potential even at E16. This sustained neurogenic activity suggests that GFAP-NPCs support cortical assembly over a longer timeframe.

The glial differentiation patterns further underscore the complementary roles of these progenitors. At E12, GFAP-NPCs produce a higher proportion of PDGFRα+ glial cells than NG2-NPCs, both at P0 and P90. However, within progeny derived from E16 NG2-NPCs begin to surpass GFAP-NPCs in glial output, reflecting a sequential transition to gliogenesis. These distinct temporal dynamics reinforce the adaptability and specialization of NG2-NPCs and GFAP-NPCs in response to the developmental needs of the cortex, probably regulated by both extrinsic and intrinsic signals (54,55). NG2-NPCs rapidly adapt to the early neurogenic requirements and later shift to gliogenesis, aligning their cellular output with the maturation demands of the cortex. GFAP-NPCs, on the other hand, initially provide a more constant supply of neurons, and transition to a gliogenic phase later, likely playing a role in stabilizing cortical assembly and glial support.

The balance between NG2+ and GFAP+ progenitors highlights the critical role of spatial and temporal coordination in cortical development and integrity. This coordination is governed by complex signaling pathways that regulate progenitor maintenance, fate commitment, and lineage progression, ultimately ensuring balanced neurogenesis and gliogenesis. Disruptions in their regulation may lead to abnormal proliferation, differentiation, or maintenance of NPCs, potentially resulting in cortical malformations (56,57) and neurodevelopmental disorders such as intellectual disability, epilepsy, and autism spectrum disorders (34,56). Understanding their distinct roles and interactions within specific cortical zones sheds light on mechanisms driving normal cortical development and the pathogenesis of these disorders. Together, NG2+ and GFAP+ progenitors uniquely shape the architecture and functional complexity of the cortex, underscoring the importance of dynamic, niche-specific cues in orchestrating progenitor behavior and lineage commitment.

### 4.3. NG2-progenitors vs RGCs: Common Genes and Specific Molecular Fingerprints

Transcriptomic analyses have significantly advanced our understanding of heterogeneity among neural progenitor cells (NPCs). Distinct clusters of NPCs have been identified during brain development in humans and mice (14–17) and across various regions of the subventricular zone (58,59). In this regard, our analysis revealed transcriptomic divergences between NG2+cells and GFAP+cells, highlighting differences in their developmental timing and their specialized roles in central nervous system development. For instance, NG2+cells are observed to follow a more restricted differentiation pathway, whereas radial glial cells (RGCs) maintain a more generalized progenitor profile. This indicates that NG2+cells at earlier stages may act as a more committed progenitor, prioritizing neurogenic support, while GFAP+cells maintain the broader potential necessary for cortical assembly.

Both, NG2+cells and GFAP+cells express characteristic neural progenitor genes (such as *Pax6* and *Sox2*), confirming their identity as progenitors (60–62). However, differences emerge in their transcriptomic signatures, reflecting their divergent developmental roles and contributions to cortical architecture. Differential gene expression (DEG) analysis from bulk-RNA sequencing displays that NG2+cells extend beyond their traditional role as oligodendrocyte precursors. During early development, NG2+cells show an enriched transcriptomic profile associated with neurogenesis and maturation-related pathways, reinforcing their neurogenic role. In contrast, GFAP+cells exhibit gene expression profiles more closely associated with the maintenance of progenitor stages. Upregulated genes in NG2+cells, such as *Odf3l1* and *Krt6a*, are involved in cytoskeletal remodeling, suggesting a role in structural adaptation critical for neuronal differentiation. For example, a transient expression of keratin in maturating neurons has been described in the rat spinal ganglion (63). Similarly, genes like *Epha6* and *Clstn1*, key players in axonal guidance and neural circuit formation, underscore the role of NG2+cells in promoting synaptic connectivity and axonal pathfinding (64,65). Additionally, *Slc13a3* facilitates neuronal metabolic demands (66), while *Atp6v0d1*, Snap47, and *NeuroD2* contribute to developmental maturation and differentiation, facilitating the transition from progenitor to neuronal identity (67–69). Upregulated genes in NG2+cells like *Nsg1* and *Nsg2*, support synaptic function and neural differentiation, further emphasizing a role in neurogenesis (70). *Lrrc4*, which promotes excitatory synapse formation (71) is also enriched in NG2+cells, strengthening their known contributions to neuronal lineages. Our findings align with previous observations that transcriptional programs specific to cortical neuron subtypes are already active in neural stem cells (NSCs) before the birth of neurons (72). This early activation of neuron-specific transcriptional profiles suggests a preparatory mechanism within NSCs that may streamline differentiation and later contribute to neuronal identity. The presence of these early transcriptional cues in NG2+cells could serve to facilitate lineage specification.

Conversely, genes associated with progenitor maintenance are significantly downregulated in NG2+cells, reflecting their progression toward a neurogenic trajectory. For example, genes such as *Folr1*, *Sostdc1*, and *Otx2*—involved in cell proliferation, BMP signaling, and early neuroectoderm specification (73–75) —show reduced expression, indicating a marked reduction in progenitor self-renewal. Moreover, genes such as *Btg2* and *Eomes* (also known as *Tbr2*), highly expressed in NSCs within the ventricular zone (VZ) and basal progenitors in the SVZ, are similarly downregulated in NG2+cells (16,76). This downregulation demonstrates that NG2+cells diverge from the renewal-driven behavior characteristic of these progenitor populations, instead favoring differentiation programs that prime them for neuronal functions. The downregulation of *Dct* further reveals a reduced proliferative potential, highlighting a decrease in proliferative potential and underscoring the commitment of NG2+cells to differentiation over stem-like maintenance (77). This unique trajectory positions NG2+cells as a specialized progenitor pool with a distinct role in neurogenesis complementing the more self-renewing capability of RGCs. Unlike RGCs, which sustain their multipotent status for extended periods, NG2+cells adopt a more lineage-restricted role, focused on supporting synaptic integration and circuit development. This functional specialization aligns with findings from gene ontology analyses, which emphasize NG2+ cells’ contributions to synaptic and structural maturation.

Among the downregulated genes in NG2+cells, several are bioinformatically predicted genes whose functions remain uncharacterized. Many of these likely represent long-non-coding RNA (lncRNAs) that are increasingly recognized as critical regulators in brain development. lncRNAs often play essential roles in neural differentiation, maintaining NSC properties, and orchestrating neurogenesis (78). Understanding the functional significance of these lncRNAs in the context of NG2+cells may clarify novel regulatory networks that guide brain development and facilitate complementary interactions between distinct NPC populations.

The downregulation of progenitor-associated genes and the enrichment of differentiation-linked pathways in NG2+cells underscore their critical role in neurogenesis and neuronal network formation. This developmental trajectory highlights the complementary functions of NG2+cells and RGCs, with NG2+cells providing a specialized contribution to neuronal lineage specification and synaptic integration. Additionally, uncharacterized genes and lncRNAs present an opportunity to unravel the intricate mechanisms regulating NPC specialization, offering new perspectives on brain development and the potential for addressing neurodevelopmental disorders linked to progenitor dysfunction.

### 4.4. Conclusion

This study highlights the distinct roles and molecular identities of NG2 and GFAP progenitors during cortical development, with each population exhibiting distinct molecular signatures, temporal dynamics, and differentiation potentials. NG2-NPCs demonstrate strong neurogenic potential, with a transcriptomic profile favoring neuron formation and synaptic establishment during early stages, transitioning to gliogenesis as development progresses. Conversely, GFAP progenitors (RGCs) exhibit a molecular profile that preserves their progenitor state during early development, showing an extended neurogenic phase before transitioning to gliogenesis, supporting their role in sustained cortical assembly. Together, these populations exhibit complementary dynamics, contributing uniquely to the balance of neurogenesis and gliogenesis. Further investigation into the signaling pathways guiding these temporal and lineage-specific transitions will provide deeper insights into CNS development mechanisms and their implications for neurodevelopmental disorders.

## 6. Supplementary Figures

**Figura S1.**
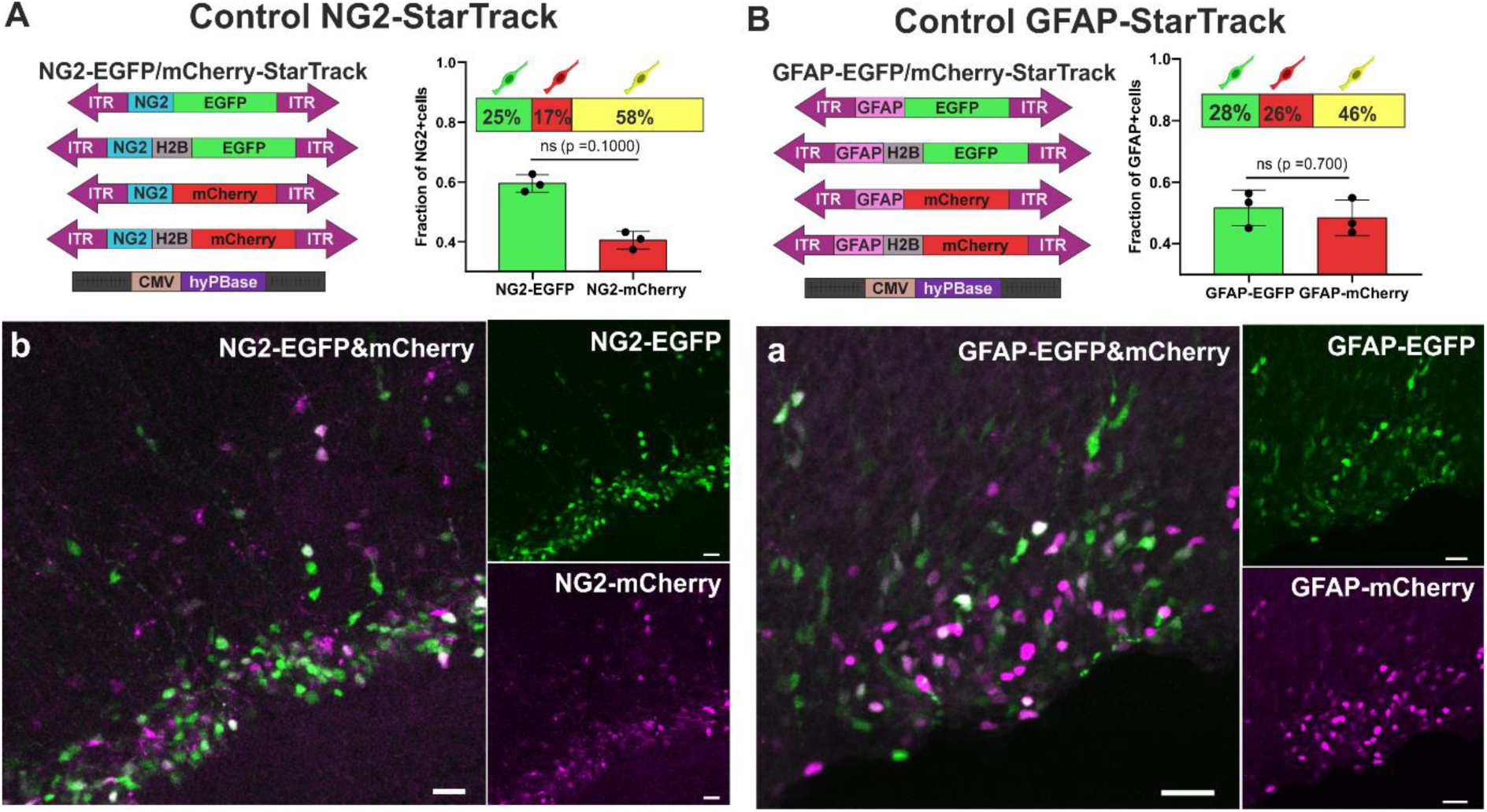
Control of plasmids coexpression. (A) StarTrack plasmid used as a control of co-expression of GFP and mcherry under the NG2-promoter. Left graph represent the percentaje of mcherry+, GFP+ or GFP&mCherry+. Barplot represent the fraction of GFP and mCherry+cells in each sample. (Ab) Image of co-electroporation under NG2-EGFP (green) and NG2-mCherry (magenta). (B) StarTrack plasmid used as a control of co-expression of GFP and mcherry under the GFAP-promoter. Left graph represent the percentaje of mcherry+, GFP+ or GFP&mCherry+. Barplot represent the fraction of GFP and mCherry+cells in each sample. (Ab) Image of co-electroporation under GFAP-EGFP (green) and GFAP-mCherry (magenta). Scale bar: 25μm.

## Notes

### Competing Interest Statement

The authors have declared no competing interest.

